# Enteral Activation of WR-2721 Mediates Radioprotection and Improved Survival from Lethal Fractionated Radiation

**DOI:** 10.1101/289074

**Authors:** Jessica M. Molkentine, Tara N. Fujimoto, Thomas D. Horvath, Aaron J. Grossberg, Amit Deorukhkar, Errol L.G. Samuel, Wai Kin Chan, Philip L. Lorenzi, Robert Dantzer, James M. Tour, Kathryn A. Mason, Cullen M. Taniguchi

**Affiliations:** Department of Experimental Radiation Oncology, The University of Texas MD Anderson Cancer Center, Houston, Texas, 77030, United States of America; Department of Bioinformatics and Computational Biology, The University of Texas MD Anderson Cancer Center, Houston, Texas, 77030, United States of America; Department of Radiation Oncology, The University of Texas MD Anderson Cancer Center, Houston, Texas, 77030, United States of America; Department of Symptoms Research, The University of Texas MD Anderson Cancer Center, Houston, Texas, United States of America; Department of Chemistry, Smalley-Curl Institute and the NanoCarbon Center, and Department of Materials Science and NanoEngineering, Rice University, Houston, Texas, United States of America

**Author notes:** Corresponding author: (CT).

## Abstract

Unresectable pancreatic cancer is almost universally lethal because chemotherapy and radiation cannot completely stop the growth of the cancer. The major problem with using radiation to approximate surgery in unresectable disease is that the radiation dose required to ablate pancreatic cancer exceeds the tolerance of the nearby duodenum. WR-2721, also known as amifostine, is a well-known radioprotector, but has significant clinical toxicities when given systemically. WR-2721 is a prodrug and is converted to its active metabolite, WR-1065, by alkaline phosphatases in normal tissues. The small intestine is highly enriched in these activating enzymes, and thus we reasoned that oral administration of WR-2721 just before radiation would result in localized production of the radioprotective WR-1065 in the small intestine, providing protective benefits without the significant systemic side effects. Here, we show that oral WR-2721 is as effective as intraperitoneal WR-2721 in promoting survival of intestinal crypt clonogens after morbid irradiation. Furthermore, oral WR-2721 confers full radioprotection and survival after lethal upper abdominal irradiation of 12.5Gy x 5 fractions (total of 62.5Gy, D2EQ=140.6Gy), which would likely ablate pancreatic cancer. We find that the efficacy of oral WR-2721 stems from its selective accumulation in the intestine, but not in tumors or other normal tissues, as determined by *in vivo* mass spectrometry analysis. Thus, we demonstrate that oral WR-2721 is a well-tolerated, and quantitatively selective, radioprotector of the intestinal tract that is capable of enabling clinically relevant ablative doses of radiation to the upper abdomen without unacceptable gastrointestinal toxicity.

## Introduction

The major limiting factor in delivering the appropriate tumoricidal dose of radiation is normal tissue toxicity to adjacent organs^1^. This issue is highlighted by solid tumors of the abdomen and pelvis, such as pancreatic and prostate adenocarcinoma, which often cannot achieve tumoricidal doses without significant morbidity to the gastrointestinal (GI) tract. For instance, pancreatic cancer often occurs in the head of the pancreas, which shares a blood supply with the duodenum, a very radiosensitive portion of the intestinal tract. Tumors of the pancreatic head require doses that exceed 77Gy to achieve local control^2^ which is often impossible to administer safely because the adjacent duodenum can only tolerate a maximum of 50Gy without causing bleeding ulcers or perforation^3,4^. Unfortunately for patients with unresectable pancreatic cancer, there are no effective treatments that specifically protect the GI tract from this radiotoxicity, and thus ablative radiotherapy is not currently possible.

WR-2721 (*S*-2-[3-aminopropylamino]-ethylphosphorothioic acid), also known as amifostine, is a proven radioprotector of normal tissues and is FDA-approved for intravenous administration. When given intravenously, amifostine causes severe nausea and hypotension^5^, which has caused the drug to fall out of favor. WR-2721 exists as a prodrug that is hydrolyzed to the active cytoprotective free thiol metabolite, WR-1065^6^ by non-specific tissue alkaline phosphatases that are enriched in normal tissues^7^. Unfortunately these ubiquitous cellular enzymes present in the endothelium may mediate undesirable side effects in the autonomic nervous system^8^.

WR-2721 was not formulated for oral delivery due to significant enteral metabolism, which would preclude therapeutic levels from accumulating in the serum and reaching the target organs such as the salivary glands^9^. The entire intestinal tract is enriched with intestinal alkaline phosphatases, which exhibit the highest levels of expression in the duodenum and jejunum^10^. Thus, we reasoned that a dose of WR-2721 given orally just before radiation could be rapidly activated by the endogenous digestive enzymes in the duodenum and jejunum to its active WR-1065 metabolite. This enterally-activated form of WR-2721 would accumulate in high concentrations in the intestines an provide localized radioprotection with potentially fewer systemic side effects. This could be particularly useful during radiation for pancreatic cancer, since the duodenum and jejunum are dose-limiting organs preventing ablative treatments.

Here, we demonstrate that oral WR-2721 is an effective radioprotector against otherwise lethal doses of radiation directed to the upper abdomen. Furthermore, we demonstrate that the drug is well–tolerated and accumulates its active metabolite, WR-1065 in significantly higher levels in the GI tract compared to the serum, liver or even spontaneous pancreatic tumors.

## Results

### Oral WR-2721 promotes intestinal crypt survival after irradiation

Systemic dosing of WR-2721 by intraperitoneal (IP) injection has already been shown to be effective at radioprotecting the gut of C3Hf/KamLaw mice receiving whole body irradiation^11,12^, but we wanted to determine if oral (PO) administration of WR-2721 would be similarly efficacious. We tested a range of doses of oral WR-2721 from 0 to 1000mg/kg, followed either 15 or 30 minutes after by a morbid dose of whole body irradiation (WBI). All mice were then subjected to a microcolony assay, which is considered the gold standard assay for demonstrating an effect of an intervention on regenerating intestinal crypts after a cytotoxic insult^13^. Indeed, compared to vehicle controls, oral WR-2721 improved crypt survival in the duodenum when administered at either 15 or 30 min prior to WBI (Figures 1A and 1B). Similarly, oral WR-2721 also promoted crypt survival in the jejunum at both 15 min (Figure 1C) and 30 min (Figure 1D) after drug treatment. Oral WR-2721 protected the duodenum and jejunum at all doses and timepoints compared to PBS controls except the lowest dose in the duodenum at 15 min. We performed a more extended dose response in the jejunum for WR-2721 given 30 min prior to 12Gy of WBI, and concluded that radiation protection was likely maximized at 500mg/kg, with no further benefit conferred by doubling the dose to 1000mg/kg (Figure 1D). There did not appear to be benefit of one time point over the other in these studies. Representative H&E stained jejunal sections from the microcolony assays are shown in figure 1E.

### Orally administered WR-2721 is better tolerated than intraperitoneal WR-2721

Severe nausea is a common and worrisome side effect of systemic WR-2721, and we were concerned that oral administration of the drug might exacerbate this side effect. Although mice do not produce an emetic response to noxious stimuli^14^, nausea is manifested as a decrease in body weight and food intake. Thus, we closely monitored body weight and food consumption as a model of nausea in mice. Based on the results of the microcolony assay (Figure 1), we used 500mg/kg as the dose for toxicity testing. Mice were treated daily for 5 consecutive days with vehicle or WR-2721 administered IP or PO, and body weight and food consumption were monitored daily (Figure 2A and 2B, respectively). There was only one vehicle control group that was subjected to both gavage and intraperitoneal injections to more stringently control for the stress of drug administration from either IP or PO route. Despite this intensive intervention, the vehicle control animals maintained their initial bodyweight and food consumption throughout the study (Figure 2A and 2B). Mice receiving intraperitoneal WR-2721 at 500mg/kg, however, exhibited a dramatic decrease in food consumption and body weight, requiring euthanasia by day 4 (Figure 2A and 2B). In contradistinction, mice receiving oral WR-2721 had a modest initial decrease in body weight and food consumption, but rapidly regained body weight with a complete recovery within 2 days of stopping treatment (Figure 2A and 2B). These data suggest that PO administration of WR-2721 is better tolerated than systemic administration and moreover, 500mg/kg orally is a feasible dose for short-term radioprotection.

### WR-2721 protects against high-dose fractionated radiation

In order to ablate pancreatic cancer with ionizing radiation, radioprotection of nearby duodenum and jejunum is required. We reasoned that orally administered WR-2721 would be directly activated in the intestine as radiation is given to maximize efficacy without systemic absorption of the drug. We modeled the transit time of oral WR-2721 with a bolus of methylene blue and assessed the progression of the dye front through the GI tract at 5-min intervals (Figure S1). We found that methylene blue reached the duodenum in 10 min and was in the jejunum by 15-30 min (Figure S1). Notably, within 30 min, the dye had not reached the cecum or large intestine. Thus, we reasoned that oral gavage of WR-2721 25 min prior to irradiation would be used in future radioprotection studies.

Stereotactic body radiotherapy (SBRT) can deliver higher and more conformal radiation doses to a smaller volume over a course of 1-5 treatments with the use of advanced image guidance. SBRT, however, is still constrained by the anatomy of the pancreas, where pancreatic tumors often abut or invade the nearby duodenum and jejunum. We designed an SBRT treatment that used 10mm radiation fields that could be used to treat pancreatic cancer in mice (Figure 3A). We correlated the radiological images with gross anatomy after methylene blue gavage to determine that even in this focused radiation field that a high dose to affect the entire pancreas, duodenum, and jejunum, along with a portion of the gastric antrum and the left lobe of the liver which are anterior to the pancreas (Figure 3B).

We performed a standard radiation dose escalation study to determine the LD50/10 (dose required to produce 50% lethality in 10 days or less) for this model of focused radiation^15^. Daily fractions ranged from 10 to 13Gy per fraction, and were given daily over 5 consecutive days. We found that 13Gy x 5 (65Gy, D2EQ=124.6Gy) led to 100% death in 10 days after treatment, and 10Gy x 5 (50Gy, D2EQ=83.3Gy) exhibited 100% survival at 30 days post-treatment (Figure 3C). All other doses showed an intermediate phenotype, with a 20% 30-day survival at 12Gy x 5 (60Gy; D2EQ=110Gy) and a 60% 30-day survival of the cohort that received 11Gy x 5 (55Gy, D2EQ=96.3Gy).

We hypothesized that the maximal benefit of radioprotection would likely occur between 12 and 13Gy at 12.5Gy x 5 fractions (62.5 Gy; D2Eq=117.2Gy). Thus, a new cohort of mice was treated with 5 daily fractions of 12.5Gy to the upper abdomen in a similar field as shown in Figure 3A, with radioprotection afforded by orally administered WR-2721 or vehicle control 25 minutes prior to each radiation treatment. Remarkably, 100% of mice that received oral WR-2721 lived beyond 30 days, while all vehicle controls died in fewer than 10 days Figure 3D, log rank p=0.0035).

### Selective enrichment of WR-1065 within intestines from oral WR-2721

We hypothesized that oral WR-2721 acts via localized conversion to WR-1065 by intestinal alkaline phosphatases in the duodenum and jejunum. This would concentrate the active metabolite WR-1065 only in the dose-limiting areas of the intestine during radiation, and possibly reduce off-target effects. To understand if this was indeed the physiologic mechanism of action, we developed a mass spectrometry assay to measure the concentrations of WR-1065, the active metabolite of WR-2721, in both serum and tissues. We first assessed the pharmacokinetics of WR-1065 appearance in the plasma and other tissues after oral gavage with 500mg/kg or IP injection with 250mg/kg of WR-2721 in C57BL/6 mice (see schema in Figure 4A). Serum and tissues were collected 25 min after gavage, which approximates the distribution of WR-1065 in tissues at the time of radiation (Figure 3C, 3D). The intraperitoneal injection of WR-2721 resulted in almost a 5-fold increase in plasma concentrations of the active metabolite WR-1065, compared to oral administration (119.1±17.3 vs 27.0±7.0, IP vs oral, p=0.001, Figure 4B). Tissues were harvested and immediately processed for metabolite collection 25 min after oral gavage or IP administration of WR-2721. We found that IP injections caused a nearly homogeneous concentration of the active metabolite WR-1065 in liver (254.3±34.1 pmol WR-1065/mg tissue), duodenum (237.7±22.7 pmol WR-1065/mg tissue), and jejunum (203.7±2.9 pmol WR-1065/mg tissue, Figure 4C). Oral WR-2721, however, showed a six-to twelve-fold enhancement of WR-1065 within duodenum (586.2±97.0 pmol WR-1065/mg tissue) and jejunum (1141±104.8 pmol WR-1065/mg tissue) compared to liver (89.0±22.4 pmol WR-1065/mg tissue), indicating the specificity of tissue protection by oral WR-2721.

We next determined the specificity of WR-1065 enrichment in intestines in a genetically engineered mouse model. We bred *Kras*^LSL/+^; *Trp53*^FL/+^;*Ptf1a*^Cre/+^(KPC) mice that develop spontaneous pancreatic cancer^16^ and backcrossed them to a C57BL/6 background over 10 generations to eliminate the confounding issue of genetic variance from inbred mice. These autochthonous tumors are thought to recapitulate the desmoplasia observed in human tumors that may make these tumors more aggressive^17^. Ideally, WR-1065 should not accumulate in tumors to be an effective clinical radioprotectant. Thus, we collected plasma, liver, duodenum, jejunum and pancreatic tumors 25 min after oral administration of 500mg/kg or IP administration of 250mg/kg WR-2721 to determine the concentrations of the WR-1065 (see schema in Figure 4D). Similar to results from wild-type C57BL/6 mice, IP injections of WR-2721 in KPC mice caused a 30-fold enrichment of WR-1065 in the serum compared to oraloral WR-2721, which did not reach statistical significance (Figure 4E, p=0.07). IP injections of WR-2721 resulted in similar concentrations of the radioprotective metabolite WR-1065 in all tissues measured (Figure 4F). The concentration of WR-1065 was 420±69.8 pmol WR-1065/mg tissue in the liver, 236.2±8.6 pmol WR-1065/mg tissue in the duodenum, 275.5±5.0 pmol WR-1065/mg tissue in the jejunum and 248±30.2 pmol WR-1065/mg tissue in pancreatic tumors.

The oral administration of WR-2721 exhibited a highly selective enrichment of WR-1065 in the intestines (Figure 4F). The concentration of WR-1065 was 200.6±23.3 pmol WR-1065/mg tissue in the duodenum and 757.7±26.8 pmol WR-1065/mg tissue in the jejunum compared to only 59.9±26.8 pmol WR-1065/mg tissue in the liver and 24.5±1.9 pmol WR-1065/mg tissue in the tumor. Thus, oral WR-2721 resulted in a 10 to 40-fold enrichment of the drug in the intestines compared to tumor (Figure 4G), while systemic administration by IP injection caused an equal distribution of radioprotective drug in both the normal tissues and tumor.

## Discussion

We demonstrate that orally administered WR-2721 is well tolerated, and radioprotects the intestinal tract from ablative doses of fractionated radiation. Here, we utilize the natural gradient of intestinal alkaline phosphatases within the duodenum and jejunum to rapidly WR-2721 to the radioprotective WR-1065 within in the gut, while limiting exposure to the rest of the body where protection is not needed, and may be counterproductive. This is most readily illustrated in our experiments with genetically engineered mice that show with a sensitive mass spectrometry technique that WR-1065 accumulates significantly higher concentrations in intestines, but not within pancreatic tumors.

The route of administration matters greatly depending on which tissues should be protected from radiation, but currently we are limited to intravenous injections of amifostine. For salivary gland protection, intravenous administration is logical since this allows the protective metabolite WR-1065 to accumulate systemically then rapidly enter these organs prior to radiation. There has never been demonstration of tumor radioprotection, but nevertheless has remained a theoretical concern of systemic administration. We demonstrate this point here showing that intraperitoneal administration causes rapid equilibration of WR-1065 across normal tissues and tumors (Figure 4). Moreover, aberrant activation of amifostine in the endothelial compartments^9^, may be partially responsible for many of the unpleasant side effects of systemic administration, such as nausea and hypotension.

Alternative routes for WR-2721 administration have been tested previously with largely negative results. For instance, endorectal infusion of amifostine was found to improve toxicity profiles after pelvic radiation in Phase I studies^18,19^, but failed to meet endpoints in a larger randomized study^20^. These may be due, in part, to reliance on the ubiquitous and non-specific alkaline phosphatases present in most cells. Moreover, the rectal mucosa has a thick mucus lining and expresses low levels of intestinal alkaline phosphatase^21^, which is required to activate WR-2721^22^.

The advantage of using oral WR-2721 is that it is already an FDA approved drug, and the pathway to repurposing this drug as an orally activated agent may be shorter than developing a new chemical variant^23^ or trying to win approval of enteric or nanoparticle formulations of WR-1065^24–26^. Thus, we view these studies as a proof of principle that WR-2721, or similar thiophosphate prodrugs, could be used to selectively target the intestine for radioprotection while limiting exposure to other normal tissues thereby decreasing the severity of the side effects as we demonstrate in this study (Figure 2).

Pancreatic cancer requires a biologically equivalent dose of more than 77Gy to have a clinical benefit^27^. Currently, this is not attainable for most pancreatic tumors unless they are in a location that is at least 1cm away from bowel^2^. In addition to the possible use of oral WR-2721 for improving the outcomes for pancreatic cancer patients, a similar strategy could be explored for others abdominal or pelvic cancers that cannot be treated definitively with radiation due to GI toxicity, such as hepatobiliary tumors, retroperitoneal sarcomas, or even metastatic disease within the abdomen.

## Materials and methods

### Mice

C57BL/6JLaw and C3Hf/KamLaw mice were purchased from the Department of Experimental Radiation Oncology’s specific pathogen free animal facility at the MD Anderson Cancer Center. *Kras* ^LSL/+^; *Trp53* ^FL/+^; *Ptf1a*^Cre/+^ mice were backcrossed to a pure C57BL/6 background over ten generations genotyped as described previously^28^. All mice were maintained in accordance with the Association for Assessment and Accreditation of Laboratory Animal Care and the Institutional Animal Care and Use Committee guidelines. Female C3Hf/KamLaw mice were used for the microcolony assay^13^ and were 12-14 weeks old at the start of treatment. Tumor-bearing KPC mice were used in the LC/MS-MS tissue analysis of WR-1065 concentrations. The remaining studies use male C57BL/6JLaw mice at 8 weeks of age at the start of all studies.

### WR-2721

WR-2721 Trihydrate (Toronto Research Chemicals Inc; North York, Ontario, Canada) was diluted in PBS (with 0.2 conversion factor to adjust for trihydrate) and was administered by IP injection at 25or250 or 500mg/kg or by oral gavage at doses from5from 150 to 1000 mg/kg in a volume of 0.1ml.

### Irradiation

For survival studies, SBRT was given for 5 consecutive days 25min after administration of vehicle or WR-2721 using the XRAD 225Cx small animal irradiator (Precision X-Ray; North Branford, CT) fitted with a conformal collimator to produce a circular 10mm diameter radiation field. Mice were anesthetized with isoflurane gas for CT imaging and irradiation. Each mouse was imaged using the XRAD 225Cx cone beam CT prior to irradiation in order to align the radiation field so that the cranial edge of the field was located 5mm below the diaphragm at the isocenter of the mouse. This field exposed the pancreas, duodenum, jejunum and liver to radiation. Radiation was administered AP/PA.

For the microcolony assay, a single dose of 12Gy whole body irradiation was administered 15min or 30min after WR-2721 treatment using a Pantak 300 x-ray unit (Pantak; East Haven, CT), with 300kVp X-rays at a dose rate of 1.84 Gy/min. Un-anesthetized mice were loosely restrained in a well ventilated 15 x 15 x 2cm Lucite box during WBI.

### Methylene blue gut assay

In order to determine the rate at which oral solutions such as WR-2721 traverse the digestive tract of 8-week old C57BL/6 mice following oral gavage, a 1% (w/v) solution of methylene blue in PBS was gavaged in a total volume of 0.1ml. Mice were then euthanized at 5, 10, 15, 25 and 30min after the dye administration and the progress of the methylene blue stain through the intestines was examined. Mice were anesthetized with isoflurane 8 min prior to euthanasia following the same procedure used prior to irradiation (8 min at 3% isoflurane and oxygen flow rate of 2 L/min) to account for any potential changes it might have on GI transit time. Methylene blue dye progress was examined in situ for comparison with the location of our standard radiation field and then the intestine was excised so that distance traveled could be determined.

### Survival studies

Mice were treated with WR-2721 or vehicle by IP injection or oral gavage followed by SBRT and monitored on a daily basis. Once mice became moribund (exhibited ruffled fur, hunched posture, persistent diarrhea and greater than 20% weight loss) they were euthanized. Time to euthanasia and percent survival were assessed.

### Microcolony assay

Viable jejunal crypts were quantified following a single dose of 12Gy WBI +/− a single oral dose of WR-2721 given at 15 min or 30 min prior to radiation using the microcolony assay^13^. Mice were euthanized 3 days and 14 hours after receiving WBI, and segments of jejunum and duodenum were resected and fixed in 10% neutral buffered formalin. Tissue was then embedded in paraffin, and four transverse slices of jejunum and duodenum were stained with hematoxylin and eosin per mouse. The number of regenerating crypts per transverse section was scored microscopically and averaged over the 4 sections per animal for each tissue type. All slides were scored by a single observer blinded to treatment group.

### LC/MS-MS analysis of WR-1065 tissue concentrations

Mice were treated with either 500mg/kg of WR-2721 by oral gavage, or 250 mg/kg by IP injection. After 17min had elapsed, the mice were anesthetized with isoflurane gas (3% isoflurane, oxygen flow rate of 2L/min). After 8 min of anesthesia, whole blood and tissues were collected. The total amount of time between treatment and tissue collection was 25 min.

While under isoflurane anesthesia, the whole blood sample was collected via cardiac puncture and was immediately transferred to a BD Microtainer® blood collection tube containing K_2_-EDTA (Becton Dickinson; Franklin Lakes, NJ). The K_2_-EDTA whole blood was then processed to plasma by centrifugation at 3,000*g* for 4 min. After processing, a volume of the plasma supernatant (typically 200 µL) was removed and immediately transferred to a fresh 1.4 mL Matrix polypropylene (PP) tube (Thermo Fisher Scientific; Waltham, MA) containing a 50 µL aliquot of 10% trichloroacetic acid (TCA). After vortexing, the TCA-treated plasma samples were stored on ice for transport from the vivarium to the laboratory for further processing. The TCA-treated plasma samples were then centrifuged at 17,000*g* for 5 min, and the supernatant was transferred to a fresh 1.4 mL Matrix tube, and the samples were either extracted and analyzed immediately or capped for −80 °C storage until analysis.

After blood collection, the mouse was euthanized, the duodenum, jejunum, liver, and tumor tissue samples were collected, and each tissue sample was placed into individually labeled PP tubes and weighed. After weighing, an aliquot of 2% TCA was added to each tube (the tissue density at 200 mg tissue per mL of solution or below for all samples), and the tissues were homogenized using a Polytron PT 1200 E hand-held homogenizer. After homogenization, the TCA-treated tissue samples were stored on ice for transport from the vivarium to the laboratory for further processing. The TCA-treated tissue samples were then centrifuged at 4,500*g* for 5 min, and the supernatant was transferred to a fresh 2 mL Simport tube (Simport; Montreal, Canada), and the samples were either extracted and analyzed immediately or capped for −80 °C storage until analysis.

### Preparation of the 2% TCA (w/v) treated C57BL/6 mouse K_2_-EDTA plasma matrix

A 2% TCA (w/v) treated plasma matrix solution was prepared by combining 0.400 mg TCA with 20 mL of C57BL/6 mouse K2-EDTA plasma (BioreclamationIVT; New York, NY) in a 50 mL Falcon PP conical tube (Thermo Fisher Scientific). The mixture was vortexed for 2 min and centrifuged at 4,500 *g* for 5 min to settle all of the precipitated protein. The 2% TCA-treated plasma matrix supernatant was transferred into two 15 mL PP tubes for long-term storage. The 2% TCA plasma matrix was maintained ice-cold while in use, and stored at −80 °C when not in use.

### Preparation of the 2% TCA (w/v) treated C57BL/6 mouse duodenal tissue matrix

A 2% TCA treated duodenal tissue matrix solution was prepared by combining individually weighed duodenal tissue samples (BioreclamationIVT) with 1.6 mL of an aqueous 2% TCA solution in hard homogenization tubes (Bertin Instruments; Bretonneux, France), and homogenized using the pre-programmed hard tissue sample setting for two cycles using the Precellys® Evolution tissue homogenizer (Bertin Instruments). Afterwards, the homogenized tissue matrix was centrifuged at 17,000*g* for 5 min to settle the tissue debris. The 2% TCA-treated duodenal tissue matrix supernatant in each tube was transferred into a 15 mL PP tube for long-term matrix storage. The 2% TCA duodenal matrix was maintained ice-cold while in use, and stored at −80 °C when not in use.

### Preparation of WR-2721 and WR-1065 stock and intermediate solutions

WR-2721 reference standard (US Pharmacopeia USP; Rockville, MD) and WR-1065 (Sigma-Aldrich; St. Louis, MO) stocks were prepared in aqueous 10 mM ammonium acetate (pH 9.2) or aqueous 2% TCA solutions, respectively. Intermediate-concentration solutions were prepared at 100 µg/mL for each compound using consistent solvents mixed 1:1 with acetonitrile.

### Preparation of WR-1065 calibration standards

Two separate sets of WR-1065 calibration standards (Calibrators) were prepared by serial dilution using either 2% TCA-treated plasma matrix or 2% TCA-treated duodenal matrix as the diluent. After the preparation of each Calibrator in each set, the standard was vortexed for a minimum of 30 s to ensure proper mixing prior to performing the subsequent dilution. The Calibrators were prepared at 1.00, 2.00, 10.0, 100, 250, 500, 900, and 1,000 ng/mL in each matrix. The Calibrators were prepared and stored individually in labeled 2 mL Simport tubes, were maintained ice-cold while in use, and stored at −80 °C when not in use.

### Extraction method for the 2% TCA-treated plasma and tissue samples

All calibrators, 2% TCA blank plasma matrix and study samples were thawed and stored on ice for the duration of the extraction procedure. A 2 µL aliquot of an ice-cold aqueous 10% TCA (w/v) solution was added to each 1.4 mL PP Matrix sample tube. A 10 µL aliquot of each Calibrator, blank sample matrix, and study sample was added to the appropriate tube. Then, a 90 µL aliquot of ice-cold acetonitrile was added to each tube, and the samples were capped and vortexed for 2 min. Finally, the samples were centrifuged at 17,000*g* for 5 min, and then the supernatant was transferred to PP injection vials for sample analysis. A typical injection volume for analysis was 10 µL.

The extraction process for the 2% TCA-treated tissue samples was similar to the plasma extraction procedure described above, with the following exception - the tissue samples do not receive a 2 µL aliquot of an ice-cold aqueous 10% TCA solution. After performing the protein precipitation and centrifugation step as described above, the tissue sample supernatant was transferred to PP injection vials for sample analysis. A typical injection volume for analysis was 10 µL.

### LC/MS-MS method

LC/MS-MS analysis was performed using an Ultimate 3000 RSLC Ultra-High Performance Liquid Chromatography (UHPLC) system coupled to a TSQ Quantiva tandem-mass spectrometer (Thermo Fisher Scientific). A zwitterionic ZIC-pHILIC (150 x 2.1mm, 5µm particle size) analytical column (MillaporeSigma; Billerica, MA) was used to achieve baseline separation of WR-2721 and WR-1065 using mobile phase A (MPA) and mobile phase B (MPB) compositions of 95/5 acetonitrile/200mM ammonium formate with 2% formic acid, and 85/10/5 water/acetonitrile/200mM ammonium formate with 2% formic acid, respectively. The chromatographic method included a column temperature of 40°C, autosampler tray chilled to 4°C, a mobile phase flow rate of 300 µL/min, and a gradient elution program specified as follows: 0-5 min, 50% MPB; 5-5.5 min, 50-80% MPB; 5.5-11 min, 80% MPB; 11-11.5 min, 80-50% MPB; 11.5-15 min, 50% MPB. The overall cycle time of the chromatographic gradient program was 15.5 minutes per sample. The blank plasma samples after the highest Calibrator (1,000 ng/mL) was free from carryover when a solution of 1:1 acetonitrile:water with 0.1% formic acid was used as the needlewash.

The TSQ Quantiva was operated in positive ion mode and had the following source parameters: source: H-ESI; source voltage: +3,200 V; sheath gas: 50; auxiliary gas: 20; sweep gas: 1; ion transfer tube temperature: 370°C; vaporizer tube temperature: 250°C. The following SRM transitions for WR-1065 were monitored: i) *m/z*: 135.1 -> 118 (quantifier), CE: 10.3 V; ii) *m/z*: 135.1 -> 58 (confirming), CE: 16 V; and iii) *m/z*: 135.1 -> 61 (confirming), CE: 22.5 V. The MS-MS system was operated in unit/unit resolution, with an RF Lens voltage of 59 V, and the dwell time of 200 ms for each SRM transition.

### Statistical analyses

Log-Rank analysis was used for survival studies and median survival was determined with 95% confidence interval (CI). Two tailed t-tests with unequal variance were used to compare PO vs. IP plasma and tissue concentrations of WR-1065. Values less than 0.05 were considered significant.

## Figure Legends

**Figure 1. Oral WR-2721 Improves Intestinal Crypt Survival after Lethal Radiation.**

Withers-Elkind microcolony assay was performed and surviving crypts per duodenal and jejunal cross-section following treatment was quantitated for the following groups: Duodenum: **(A)** WR-2721 given PO 15 min prior (n=6-20/group) or **(B)** 30min n=6-20/group) prior to 12Gy WBI. Jejunum: **(C)** WR-2721 given PO 15 min prior (n=6-20/group) or **(D)** 30 min n=6-42/group) prior to 12Gy WBI. **(E)** Representative H&E used in **(C)** and **(D)** with arrows highlighting surviving crypts.

**Figure 2. Oral Administration of WR-2721 is Better Tolerated than Systemic Injections.**

**(A)** Body weight and **(B)** food intake was measured daily following administration of WR-2721 for 5 consecutive days by either oral gavage (green triangles, n=6), IP injection (red squares, n=5), or vehicle control (blue circles, n=5).

**Figure 3. Oral WR-2721 improves survival after lethal fractionated radiation**

**(A)** Cone beam CT of a mouse taken just before irradiation with an overlay of where the 10mm radiation field would be located. Tick marks within the reticle denote 5mm intervals. **(B)** Mouse gavaged with methylene blue then dissected 25 min later. Blue dye is evident within the jejunum (green arrow), but not distal intestine (white arrow) at what would be the time of irradiation following a dose of WR-2721. **(C)** Mice were treated with 5 fractions of radiation given over 5 consecutive days using a 10mm diameter circular radiation field (n=5/group). **(D)** Mice were treated with 500mg/kg of WR-2721 by oral gavage 25 min prior to each fraction of 12.5Gy for 5 consecutive days using a 10mm diameter circular radiation field (n=5/group).

**Figure 4. Selective enrichment of the radioprotective metabolite WR-1065 by oral gavage**

**(A)** Schema of C57BL/6 mice treated with WR-2721 at 500mg/kg PO or 250mg/kg IP at 25 min prior to blood and tissue collection for LC/MS-MS analysis. C57BL/6 plasma concentrations **(B)** and C57BL/6 duodenum, jejunum and liver concentrations **(C)**. Schema for WR-1065 determination in KPC animals shown in **(D)**. **(E)** KPC plasma concentrations and **(F)** KPC duodenum, jejunum, liver and pancreatic tumor concentrations after oral and IP injections. **(G-H)** Data from (**F**) reformatted to show as ratio of WR-1065 in the noted tissue compared to spontaneous tumors.

## Acknowledgments

**Funding:** C.M.T. was supported by funding from Cancer Prevention & Research Institute of Texas (CPRIT) grant RR140012, V Foundation (V2015-22), Kimmel Foundation, Sabin Family Foundation Fellowship and the McNair Foundation.

Supported by the NIH/NCI under award number P30CA016672 and used the Small Animal Imaging Facility.

This work was funded by Cancer Prevention Research Institute of Texas grant RP130397.

## Author contributions

J.M.M. and T.N.F. executed most experiments. J.M.M. performed a primary analysis for most experiments. T.D.H. developed the LC/MS-MS assay under the supervision of P.L.L. J.M.M., T.D.H., W.K.C., A.D. and T.N.F. harvested tissues for LC/MS-MS analysis. J.M.M., C.M.T., K.A.M., J.M.T. and A.J.G. designed experiments. The manuscript was written by J.M.M., C.M.T. and T.D.H.

